# Deep Learning–Based Reconstruction: Model Comparison for Variable-Density GRAPPA ^1^H MRSI

**DOI:** 10.64898/2026.06.17.733011

**Authors:** Xinyu Zhang, Mahrshi Jani, Andrew Wright, Kimberly L. Chan, Anke Henning

## Abstract

Proton magnetic resonance spectroscopic imaging (^1^H MRSI) enables quantitative mapping of brain metabolites, but its clinical use remains limited by long acquisition time. The goal of this work to improve the applicability of high-resolution ^1^H FID-MRSI at 7T by enhancing GRAPPA-based acceleration through deep learning–driven k-space reconstruction. In particular, compared with conventional GRAPPA, MultiNet PyGRAPPA enables substantially higher in-plane acceleration while suppressing residual lipid aliasing and preserving metabolite map fidelity in non–lipid-suppressed MRSI. Building on the MultiNet PyGRAPPA framework, we introduce a comprehensive comparison of advanced machine-learning models for predicting missing k-space points. Multiple architectures—including multilayer perceptrons, convolutional neural networks, and several U-Net variants—were trained within a variable-density k-space undersampling scheme to support acceleration factors of R = 4, 6, and 7. The proposed U-Net model extends the MultiNet concept by leveraging nonlinear hierarchical feature extraction, thereby improving reconstruction fidelity while maintaining robustness to noise.The methods were evaluated in vivo using retrospectively undersampled 7T ^1^H FID-MRSI datasets from healthy volunteers and patients. Quantitative analyses demonstrate that the U-Net outperforms the original MultiNet approach, offering improved SNR retention rate, reduced lipid RMSE, and higher structural similarity of major metabolites. Metabolite maps reconstructed with the U-Net showed reduced lipid artifacts and improved anatomical consistency. In conclusion, integrating deep convolutional networks into GRAPPA-based k-space prediction provides a more reliable and higher-fidelity reconstruction pipeline. When combined with variable-density undersampling, this approach enables faster acquisition of high-resolution ^1^H MRSI without compromising spectral quality or metabolite quantification.

## 1. INTRODUCTION

Proton (^1^H) Magnetic Resonance Spectroscopy (MRS) is a non-invasive technique that enables the detection and quantification of brain metabolites, with many clinical and research applications in neurological disorders such as brain tumors and epilepsy (Rudkin et al., 1999; Cebeci et al., 2022; Maudsley et al., 2021; Wilson et al., 2019). Despite its potential, the clinical utility of proton Magnetic Resonance Spectroscopic Imaging (^1^H MRSI) remains constrained by prolonged acquisition times and its susceptibility to motion artifacts.

To mitigate these challenges, faster acquisition strategies like k-space undersampling schemes including parallel imaging, compressed sensing and multi-band have been employed to expedite MRSI scans (Bogner et al., 2021). Among these, GRAPPA, a parallel imaging technique, has enhanced imaging speed without requiring explicit knowledge of coil sensitivity profiles, thereby offering greater flexibility across various phase-encoding directions (Griswold et al., 2002). Furthermore, because GRAPPA does not rely on explicit sensitivity map estimation, it is generally more robust to errors in coil sensitivity profiles, whereas inaccuracies in these maps can lead to elevated aliasing artifacts in SENSE-like techniques (Bauer et al., 2007). In addition, the implementation of variable density sampling—where acceleration is more pronounced in the outer k-space regions than in the center—has been shown to achieve higher acceleration factors without compromising image quality, due to the oversampling of central k-space that mitigates high-energy foldover artifacts (Peters et al., 2000).

In parallel with advances in acquisition techniques, the rapid development of deep learning has spurred significant interest in applying neural network-based methods to Magnetic Resonance Imaging (MRI) and Magnetic Resonance Spectroscopy Imaging (MRSI). Early studies in MRI demonstrated that neural networks could effectively mitigate artifacts arising from k-space undersampling (Yan et al., 1993). Building on this foundation, more recent approaches have leveraged sophisticated deep learning architectures including convolutional neural networks (CNNs) and U-Nets, to reconstruct highly accelerated MRI data (Schlemper et al., 2018; Souza et al., 2019; Wang et al., 2020). Furthermore, hybrid models that integrate residual connections and attention mechanisms have been developed to enhance reconstructions in both the frequency and image domains, substantially reducing aliasing and noise artifacts (Hossain et al., 2023; Huang et al., 2018; Lan et al., 2023).

In the realm of Magnetic Resonance Spectroscopic Imaging (MRSI), most efforts of deep learning applications have concentrated on processing and analysis (van de Sande et al., 2023). However, relatively few studies have explored reconstruction methods. Neural networks (NNs) were employed to predict missing k-space data in GRAPPA-based reconstructions, improving signal-to-noise ratio (SNR) and reducing aliasing artifacts through subject-specific or semi-synthetic calibration strategies (Nassirpour et al., 2018; Chan et al., 2022). U-Net architectures emerged as a prominent approach, with one framework operating in the frequency domain to suppress truncation artifacts in truncated free induction decay (FID) signals, while another effectively reconstructed non-uniformly sampled 2D L-COSY spectra, outperforming traditional compressed sensing techniques (Lee et al., 2020; Iqbal et al., 2021). For multi-dimensional spectroscopy, an encoder-decoder HRNet demonstrated superior reconstruction of fully sampled L-COSY spectra from sparse inputs compared to iterative methods (Luo et al., 2020). Geometric deep learning was explored via shallow graph convolutional neural networks (CNNs) to optimize k-space-based coil combination, enhancing robustness across SNR variations (Motyka et al., 2021). In addition, a deep convolutional neural network (CNN)–based framework, Deep-ER, was developed for ECCENTRIC spectroscopic imaging, leveraging joint k-space and image-space feature learning with residual connections to enable fast, high-resolution reconstruction while maintaining spectral fidelity (Weiser et al., 2025).

These advancements underscore the integration of convolutional architectures and hybrid physics-guided AI strategies, driving improvements in artifact suppression, accelerated acquisition, and spectral accuracy in MRSI. Alongside deep learning approaches, physics-based subspace modeling has also contributed to reconstruction advancements. The SPICE (SPectroscopic Imaging by exploiting spatiotemporal CorrElation) method, reconstructs MRSI data within a low-dimensional spectral–temporal subspace using a hybrid CSI–EPSI acquisition (Lam et al., 2016). This method enables high-resolution 2D/3D metabolite mapping with strong artifact suppression and SNR preservation. However, because SPICE operates entirely in the spectral–temporal domain rather than directly reconstructing undersampled k-space data, it addresses a different reconstruction problem and does not target accelerated k-space acquisition.

In this study, we build upon the foundational Multi-Net GRAPPA method, which employed a simple Multi-Layer Perceptron (MLP) optimized with the Broy-den–Fletcher–Goldfarb–Shanno (BFGS) algorithm (Nassirpour et al., 2018). Additionally, we adapt a semi-synthetic calibration dataset generation technique that has been shown to enhance signal-to-noise performance (Chan et al., 2022). To further enhance reconstruction performance, we systematically evaluate more advanced neural network architectures, including MLPs with stochastic gradient descent (SGD) and Adam optimizers, Convolutional Neural Network (CNN) and U-Nets with varying depths including VGG16 U-Net, U-Net++, Attention U-Net and Residual U-Net (Robbins et al., 1951; Kingma et al., 2014; LeCun et al., 1998; Ronneberger et al., 2015; Simonyan et al., 2014; Zhou et al., 2018; Oktay et al., 2018; He et al., 2016; Kugelman et al., 2022). Our results demonstrate that the top-performing model, a U-Net, enables high-resolution 7T brain metabolite mapping while substantially enhancing SNR retention and reducing lipid contamination compared to the original MLP approach. Across multiple acceleration factors (R = 4, 6, 7), the U-Net consistently improves median SNR retention rate by approximately 16–23% and reduces lipid RMS by 5–10%, highlighting its ability to maintain spectral fidelity and artifact suppression. These improvements underscore the potential of deep learning-based k-space reconstruction to accelerate MRSI acquisition without compromising data quality.

## 2 Methods

### 2.1 In vivo data

Five healthy volunteers (three male, two female, age: 31.2 ± 7.3 y) and three patients with brain tumors (one male, two female, age = 51.3 ± 30.1 y) were enrolled after providing written informed consent prior to the measurements. We have 3 patients with low grade gliomas before therapy.

Elliptically sampled ^1^H MRSI data were acquired on an Philips 7T DSync human MRI scanner with a dual-transmit and 32-channel receive head coil (Nova Medical). A short-TE and TR 2D ^1^H FID-MRSI sequence without lipid suppression (Henning et al., 2009) was used with an optimized water suppression scheme. Data were acquired with a flip angle of 33°, a TR/TE of 322/1.21 ms, an acquisition delay of 1.3 ms, a spectral band-width of 4000 Hz, 1024 spectral points, and total acquisition duration of 11 min. All data had a slice thickness of 12 mm and used a 50 × 50 phase encoding matrix yielding a voxel size of 4.4 mm × 4.4 mm. Prior to MRSI data acquisition, B_0_ field homogeneity was optimized using field-map-based 2nd-order shimming with the custom-developed OmniShim tool (Jani et al., 2025). A high-resolution anatomical image 5 × 5 times the resolution of the MRSI data was also acquired using a T1w-FFE sequence, a flip angle of 12° and a TR of 55 ms at the same position with the same slice thickness and field-of-view as the MRSI data and a voxel size of 0.7 × 0.7 × 12 mm^3^ in a total acquisition time of 16 s. All data were exported in a raw data format so that complex coil data with both magnitude and phase information were available.

The ^1^H FID MRSI data were retrospectively undersampled by a factor of 4 and reconstructed using deep learning models trained on anatomical and auto-calibration data, as previously described (Nassirpour et al., 2018; Chan et al., 2022). Retrospective lipid removal with L2 regularization was also applied to the reconstructed data (Bilgic et al., 2013). Data were post-processed as previously described with eddy current correction, automatic phase correction, residual water removal, coil combination, and missing point prediction (Chan et al., 2022). Metabolite maps were generated by fitting the processed data with LCModel (Provencher et al., 1993) from 0.6–4.2 ppm with a basis set simulated using VESPA (Soher et al., 2011). This basis set included glycerophosphocholine, taurine, creatine, glucose, N-acetylaspartate, lactate, GABA, N-acetylaspartylglutamate, glutathione, glutamate, myo-inositol, phosphocholine, glutamine, scyllo-inositol, and aspartate with a linewidth of 3 Hz. Relaxation-corrected macromolecules were simulated and integrated into the basis set as described previously (Wright et al., 2021), and residual baseline distortions were modeled by a spline baseline (dkntmn) parameter of 0.25 in LCModel. Metabolite maps were corrected for T1 relaxation effects and displayed as described previously (Wright et al., 2022).

### 2.2 Deep learning models

The original MultiNet GRAPPA utilizes a sequential prediction strategy that we preserve in this work: missing k-space points are reconstructed through an iterative process wherein a cross-neighbor prediction is first applied, followed by an adjacent-neighbor prediction, with this two-step procedure repeated cyclically until the entirety of k-space is filled (Nassirpour et al., 2018). In the original implementation, MLP weights were optimized using the L-BFGS algorithm; in the present study, we additionally evaluate stochastic gradient descent (SGD) and Adam on the same MLP architecture (Robbins et al., 1951; Kingma et al., 2014; LeCun et al., 1998). Subsequently, a convolutional neural network was trained using kernel sizes of 2 or 3 and various activation functions (linear, sigmoid, and ReLU) to determine the most effective configuration for the prediction task. We advanced reconstruction performance through a series of stepwise improvements employing U-Nets with different depth and variants (Ronneberger et al., 2015). In particular, VGG16 U-Net leverages the pre-trained VGG16 network as its encoder, benefiting from rich hierarchical feature representations learned from large-scale datasets, which enhances the model’s ability to capture spatial and contextual information for improved reconstruction performance (Simonyan et al., 2014).U-Net++ integrates dense skip connections to more effectively bridge the semantic gap between encoder and decoder features, thereby enhancing feature aggregation and gradient flow (Zhou et al., 2018). The Attention U-Net introduces attention gates within its skip connections to emphasize salient spatial regions and suppress extraneous background information, which contributes to improved localization and overall reconstruction accuracy (Oktay et al., 2018). Finally, the Residual U-Net incorporates residual learning via shortcut connections in each layer, facilitating the optimization of residual mappings and further promoting efficient gradient propagation during training (He et al., 2016). A visual summary of each architecture is given in Figure 1. All experiments are undertaken using Tensorflow 2.13.1 in Python 3.8 on the UTSW BioHPC cluster (CPU node with 512 GB RAM). A consistent training protocol was applied across all models, incorporating early stopping and a one-cycle learning rate policy to prevent overfitting and accelerate convergence.

**Figure 1.**
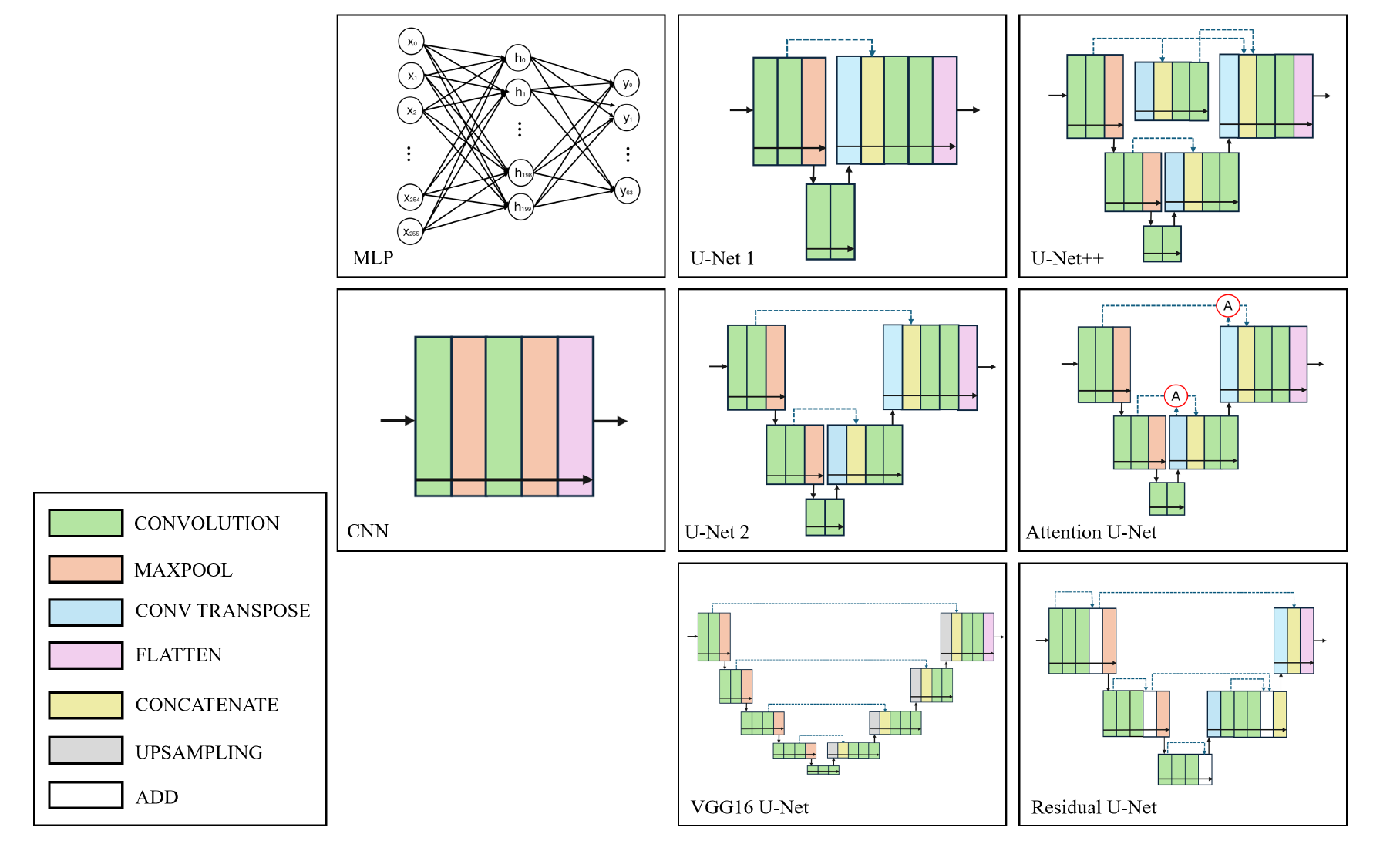
Summary of the deep learning architectures compared in this study.

### 2.3 Variable-density acceleration schemes

In previous MultiNet implementations using a 9.4T human MRI scanner (Siemens Magnetom) and a 7T human MRI scanner (Philips Dsync), variable-density acceleration schemes were introduced to achieve 4×, 6× and 7× acceleration, incorporating Philips’ elliptical k-space sampling strategy (Nassirpour et al., 2018; Chan et al., 2022). Herein initially, all models were trained using the 4× acceleration rate to facilitate direct comparisons. For the best-performing model, additional acceleration schemes (6× and 7×) were subsequently applied to further exploit and enhance its reconstruction performance.

### 2.4 Comparison metrics

The reconstruction models were evaluated using multiple quantitative metrics, including signal-to-noise ratio (SNR), root-mean-square errors (RMSEs) for lipid and five major metabolites (NAA, Cr, Cho, mI, and Glu), and corresponding Cramér-Rao lower bounds (CRLBs). SNR was calculated as the ratio between the Cr peak (2.8-3.2 ppm) and the root-mean-square (RMS) noise (10-12 ppm).

Lipid RMSE was calculated as the root-mean-square of the lipid signal (0.3–1.8 ppm). The RMSE of the metabolite maps is calculated for each metabolite by:

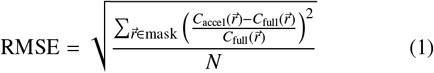

where *C* is the concentration of the metabolite and *N* is the number of voxels in the spatial mask. The CRLBs are taken from the values reported from LCModel (Provencher et al., 1993). The structural similarity index (SSIM) was used to compare the structural similarity of the metabolite maps between the fully elliptically-sampled data and the accelerated data (Wang et al., 2004). This metric is defined as:

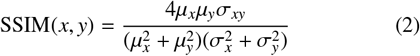

where *x* and *y* are the two signals being compared, µ_*x*_ and µ_*y*_ are the corresponding means, σ_*x*_ and σ_*y*_ are the corresponding standard deviations, and σ_*xy*_ is the correlation coefficient. All metrics were calculated across the eight subjects that shared the same acquisition parameters.

### 2.5 Model comparison

To identify the optimal reconstruction model, we employed a rank-based composite scoring approach across all 17 evaluation metrics (Cr SNR, lipid RMSE, NAA RMSE, Cr RMSE, Cho RMSE, mI RMSE, Glu RMSE, NAA CRLB, Cr CRLB, Cho CRLB, mI CRLB, Glu CRLB, NAA SSIM, Cr SSIM, Cho SSIM, mI SSIM, Glu SSIM). For each metric, the 13 models were independently ranked from 1 (worst) to 13 (best). Metrics in which higher values indicate better performance (SNR and SSIM) were ranked in ascending order, while those in which lower values denote better performance (RMSE and CRLB) were ranked in descending order.

Each model’s total performance score was obtained by summing its ranks across the 17 metrics. The maximum possible score corresponds to a model ranked best across all metrics (221), while the minimum corresponds to the lowest rank across all metrics (17). To enable comparison across models, scores were normalized to a percentage scale, with 100% representing the best possible overall performance.

## 3. Results

Table 1 and Table 2 present the quantitative results of various deep learning models for 4-fold accelerated ^1^H MRSI re-construction across five healthy volunteers and three patient datasets. Compared to the original MLP model utilizing the LBFGS solver, MLP with SGD and Adam solvers, Attention U-Net, and Residual U-Net provided a statistically significant improvement in SNR (*p* < 0.05), while U-Net1, U-Net++, and U-Net2 models achieved a highly significant improvement (*p* < 0.01), with an 8.09%, 12.8%, 15.88%, 16.08%, 19.68%, 23.16%, and 24.56% increase for MLP with SGD solver, MLP with Adam solver, Attention U-Net, Residual U-Net, U-Net1, U-Net++, and U-Net2, respectively.

**Table 1.** Quantitative evaluation of the different ^1^H MRSI reconstruction methods for an acceleration factor of R = 4. Seven metrics are used for the evaluation (SNR, lipid RMSE and CRLBs (%) of the maps for five major brain metabolites). Each value represents the average across five healthy volunteers and three patients. The results of the elliptically sampled case are also shown in the first row for comparison.

| Network architecture | Solver | Kernel size | Activation | SNR | Lipid RMSE | CRLB NAA | CRLB Cr | CRLB Cho | CRLB ml | CRLB Glu |
| --- | --- | --- | --- | --- | --- | --- | --- | --- | --- | --- |
| Elliptically sampled | — | — | — | $17.56 \pm 6.25$ | — | $6.67 \pm 2.03$ | $12.28 \pm 3.40$ | $5.84 \pm 1.64$ | $11.46 \pm 2.77$ | $9.24 \pm 2.18$ |
| MLP | lbfgs | — | Linear | $11.96 \pm 4.25$ | 1.79 | $10.22 \pm 3.13$ | $13.64 \pm 4.13$ | $8.51 \pm 2.35$ | $14.71 \pm 2.79$ | $13.32 \pm 3.09$ |
| | adam | — | Linear | $13.93 \pm 6.04$ | 1.71 | $8.33 \pm 3.59$ | $14.67 \pm 5.71$ | $6.94 \pm 2.45$ | $13.85 \pm 4.58$ | $11.12 \pm 3.57$ |
| | sgd | — | Linear | $13.34 \pm 4.70$ | 1.76 | $9.43 \pm 3.44$ | $15.21 \pm 6.27$ | $7.68 \pm 2.26$ | $14.46 \pm 3.86$ | $12.38 \pm 3.16$ |
| CNN | adam | 2 | ReLU | $5.25 \pm 2.70$ | 2.14 | $9.93 \pm 7.08$ | $18.56 \pm 12.04$ | $7.70 \pm 5.16$ | $16.11 \pm 10.42$ | $13.00 \pm 7.55$ |
| | adam | 2 | Linear | $8.55 \pm 4.23$ | 1.84 | $8.35 \pm 3.95$ | $14.99 \pm 7.00$ | $6.98 \pm 2.79$ | $13.86 \pm 5.75$ | $11.37 \pm 3.62$ |
| | adam | 2 | Sigmoid | $1.91 \pm 1.07$ | 4.01 | $19.46 \pm 15.69$ | $27.00 \pm 20.73$ | $16.39 \pm 13.61$ | $22.57 \pm 16.91$ | $23.65 \pm 16.62$ |
| | adam | 3 | Linear | $9.04 \pm 4.35$ | 1.81 | $8.25 \pm 4.00$ | $15.03 \pm 7.25$ | $6.84 \pm 2.49$ | $13.53 \pm 5.23$ | $11.04 \pm 3.56$ |
| U-Net1 | adam | 3 | Linear | $14.71 \pm 5.60$ | 1.70 | $7.33 \pm 2.34$ | $13.44 \pm 5.07$ | $6.23 \pm 1.82$ | $12.50 \pm 3.39$ | $9.97 \pm 2.33$ |
| U-Net2 | adam | 3 | Linear | $15.29 \pm 5.70$ | 1.70 | $7.26 \pm 2.26$ | $12.77 \pm 3.71$ | $6.20 \pm 1.77$ | $12.22 \pm 2.91$ | $9.90 \pm 2.25$ |
| VGG16 U-Net | adam | 3 | Linear | $12.85 \pm 7.22$ | 1.98 | $10.24 \pm 8.70$ | $18.33 \pm 10.26$ | $8.78 \pm 6.35$ | $18.66 \pm 11.61$ | $16.17 \pm 11.45$ |
| U-Net++ | adam | 3 | Linear | $15.14 \pm 5.84$ | 1.70 | $7.33 \pm 2.37$ | $13.14 \pm 4.44$ | $6.22 \pm 1.77$ | $12.31 \pm 3.17$ | $9.96 \pm 2.35$ |
| Attention U-Net | adam | 3 | Linear | $14.24 \pm 6.20$ | 1.71 | $7.46 \pm 2.65$ | $13.40 \pm 4.46$ | $6.27 \pm 2.02$ | $12.62 \pm 3.27$ | $10.11 \pm 2.46$ |
| Residual U-Net | adam | 3 | Linear | $14.20 \pm 5.88$ | 1.70 | $7.47 \pm 2.61$ | $13.53 \pm 5.01$ | $6.26 \pm 1.87$ | $12.49 \pm 3.44$ | $10.06 \pm 2.45$ |

**Table 2.** Quantitative evaluation of the different ^1^H MRSI reconstruction methods for an acceleration factor of R = 4. Ten metrics are used for the evaluation (RMSEs and SSIMs of the maps for five major brain metabolites). Each value represents the average across five healthy volunteers and three patients.

| Network architecture | Solver | Kernel size | Activation | NAA RMSE | Cr RMSE | Cho RMSE | ml RMSE | Glu RMSE | SSIM NAA | SSIM Cr | SSIM Cho | SSIM ml | SSIM Glu |
| --- | --- | --- | --- | --- | --- | --- | --- | --- | --- | --- | --- | --- | --- |
| MLP | lbfgs | — | Linear | 0.18 | 0.30 | 0.15 | 0.29 | 0.19 | 0.8848 | 0.6317 | 0.8690 | 0.8331 | 0.8589 |
|  | adam | — | Linear | 0.17 | 1.17 | 0.12 | 0.33 | 0.17 | 0.8677 | 0.6834 | 0.8306 | 0.7473 | 0.8280 |
|  | sgd | — | Linear | 0.16 | 0.74 | 0.09 | 0.24 | 0.16 | 0.9038 | 0.7016 | 0.9032 | 0.8551 | 0.9076 |
| CNN | adam | 2 | ReLU | 0.33 | 0.41 | 0.25 | 0.57 | 0.37 | 0.5898 | 0.2674 | 0.4585 | 0.4236 | 0.4972 |
|  | adam | 2 | Linear | 0.21 | 0.29 | 0.17 | 0.34 | 0.23 | 0.8246 | 0.5929 | 0.7649 | 0.7442 | 0.7926 |
|  | adam | 2 | Sigmoid | 1.04 | 3.34 | 0.85 | 1.87 | 1.18 | 0.0472 | 0.0109 | 0.0175 | 0.0011 | 0.0350 |
|  | adam | 3 | Linear | 0.21 | 1.18 | 0.16 | 0.32 | 0.22 | 0.8275 | 0.6373 | 0.7981 | 0.7506 | 0.8108 |
| U-Net1 | adam | 3 | Linear | 0.13 | 0.18 | 0.12 | 0.20 | 0.13 | 0.9276 | 0.8031 | 0.9194 | 0.8980 | 0.9296 |
| U-Net2 | adam | 3 | Linear | 0.12 | 0.18 | 0.11 | 0.19 | 0.12 | 0.9449 | 0.8063 | 0.9327 | 0.9140 | 0.9385 |
| VGG16 U-Net | adam | 3 | Linear | 0.44 | 0.43 | 0.32 | 0.54 | 0.46 | 0.3756 | 0.2482 | 0.4474 | 0.4784 | 0.3446 |
| U-Net++ | adam | 3 | Linear | 0.13 | 0.24 | 0.07 | 0.22 | 0.13 | 0.9313 | 0.8097 | 0.9272 | 0.8971 | 0.9313 |
| Attention U-Net | adam | 3 | Linear | 0.14 | 0.19 | 0.11 | 0.23 | 0.15 | 0.9232 | 0.7944 | 0.9169 | 0.8854 | 0.9187 |
| Residual U-Net | adam | 3 | Linear | 0.13 | 0.20 | 0.08 | 0.23 | 0.14 | 0.9218 | 0.7854 | 0.9188 | 0.8857 | 0.9238 |

Furthermore, U-Net++, U-Net2, and U-Net1 reduced lipid RMSE by 5.12%, 5.18%, and 5.26% (*p* < 0.05) relative to the original MLP method, indicating enhanced lipid suppression. Across all models, lipid RMSE was lowest for U-Net1, U-Net2, U-Net++, and Residual U-Net. Regarding the evaluation of major metabolites—including creatine (Cr), N-acetylaspartate (NAA), choline (Cho), the combined glutamate and glutamine signal (Glx), and myo-inositol (mI)—the U-Net models, excluding VGG16 U-Net, demonstrated substantial improvements in RMSE, CRLB, and SSIM compared with the original MLP results.

Based on the rank-based composite evaluation, U-Net2 achieved the highest score, making it the best-performing model, with 95.9% of the maximum performance. Other models that outperformed the baseline MLP with L-BFGS solver (43.9%) included U-Net++ (88.2%), U-Net1 (82.4%), Residual U-Net (76.0%), Attention U-Net (71.9%), and MLP with Adam (49.8%) and SGD (52.9%) optimizers.

Table 3 presents a quantitative evaluation of different acceleration rates for the U-Net2 and the original MLP models. Among the three acceleration rates, the 4× acceleration rate demonstrated the highest SNR retention for both models. Specifically, the median SNR retention percentages for the original MLP model were 71%, 63%, and 58% at 4×, 6×, and 7× acceleration, respectively. In comparison, U-Net2 retained 87%, 74%, and 67% at the same acceleration rates. This showed that U-Net2 outperformed the original MLP-based MultiNet in maintaining SNR across all acceleration rates. In terms of overall reconstruction quality, U-Net2 achieved statistically significant improvements across all evaluated metrics at 4× acceleration (all p < 0.05), including SNR, lipid RMSE, metabolite RMSEs, CRLBs, and SSIMs. At higher acceleration factors, U-Net2 continued to outperform the original MLP on the majority of measures, with statistically significant improvements observed in 13 of 17 metrics at 6× acceleration and 7 of 17 metrics at 7× acceleration, while the remaining metrics showed comparable performance.

**Table 3.**
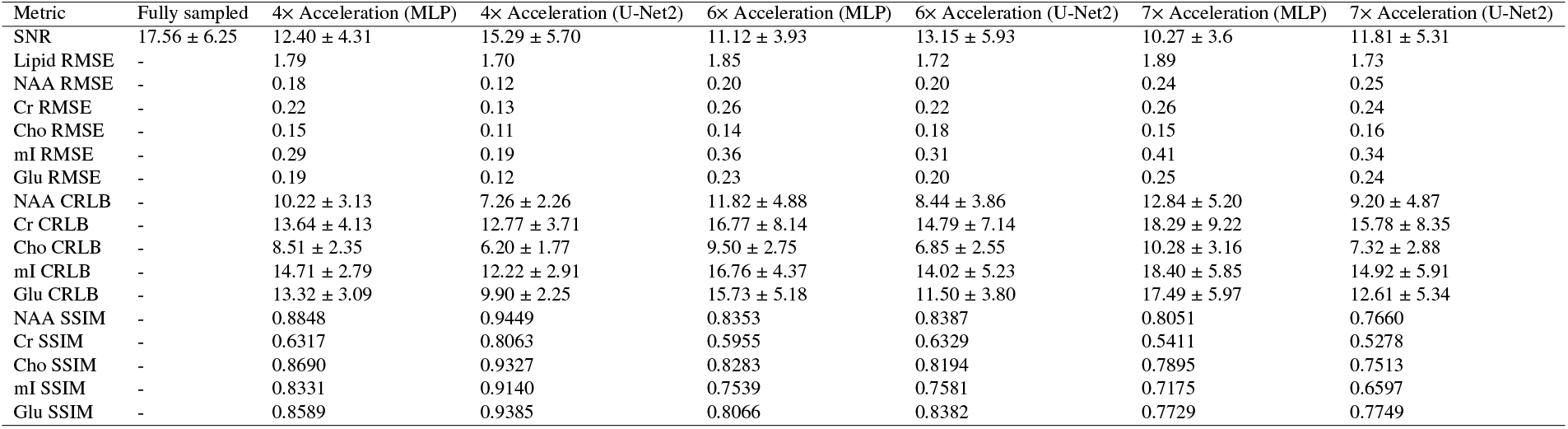
Quantitative evaluation of the different ^1^H MRSI acceleration rates for U-Net2 and the original MLP MultiNet model. Seventeen metrics are used for the evaluation (SNR, lipid RMSE, RMSEs, CRLBs (%) and SSIMs of the maps for five major brain metabolites). Each value represents the average across five healthy volunteers and three patients.

| Metric | Fully sampled | 4× Acceleration (MLP) | 4× Acceleration (U-Net2) | 6× Acceleration (MLP) | 6× Acceleration (U-Net2) | 7× Acceleration (MLP) | 7× Acceleration (U-Net2) |
| --- | --- | --- | --- | --- | --- | --- | --- |
| SNR | 17.56 ± 6.25 | 12.40 ± 4.31 | 15.29 ± 5.70 | 11.12 ± 3.93 | 13.15 ± 5.93 | 10.27 ± 3.6 | 11.81 ± 5.31 |
| Lipid RMSE | - | 1.79 | 1.70 | 1.85 | 1.72 | 1.89 | 1.73 |
| NAA RMSE | - | 0.18 | 0.12 | 0.20 | 0.20 | 0.24 | 0.25 |
| Cr RMSE | - | 0.22 | 0.13 | 0.26 | 0.22 | 0.26 | 0.24 |
| Cho RMSE | - | 0.15 | 0.11 | 0.14 | 0.18 | 0.15 | 0.16 |
| mI RMSE | - | 0.29 | 0.19 | 0.36 | 0.31 | 0.41 | 0.34 |
| Glu RMSE | - | 0.19 | 0.12 | 0.23 | 0.20 | 0.25 | 0.24 |
| NAA CRLB | - | 10.22 ± 3.13 | 7.26 ± 2.26 | 11.82 ± 4.88 | 8.44 ± 3.86 | 12.84 ± 5.20 | 9.20 ± 4.87 |
| Cr CRLB | - | 13.64 ± 4.13 | 12.77 ± 3.71 | 16.77 ± 8.14 | 14.79 ± 7.14 | 18.29 ± 9.22 | 15.78 ± 8.35 |
| Cho CRLB | - | 8.51 ± 2.35 | 6.20 ± 1.77 | 9.50 ± 2.75 | 6.85 ± 2.55 | 10.28 ± 3.16 | 7.32 ± 2.88 |
| mI CRLB | - | 14.71 ± 2.79 | 12.22 ± 2.91 | 16.76 ± 4.37 | 14.02 ± 5.23 | 18.40 ± 5.85 | 14.92 ± 5.91 |
| Glu CRLB | - | 13.32 ± 3.09 | 9.90 ± 2.25 | 15.73 ± 5.18 | 11.50 ± 3.80 | 17.49 ± 5.97 | 12.61 ± 5.34 |
| NAA SSIM | - | 0.8848 | 0.9449 | 0.8353 | 0.8387 | 0.8051 | 0.7660 |
| Cr SSIM | - | 0.6317 | 0.8063 | 0.5955 | 0.6329 | 0.5411 | 0.5278 |
| Cho SSIM | - | 0.8690 | 0.9327 | 0.8283 | 0.8194 | 0.7895 | 0.7513 |
| mI SSIM | - | 0.8331 | 0.9140 | 0.7539 | 0.7581 | 0.7175 | 0.6597 |
| Glu SSIM | - | 0.8589 | 0.9385 | 0.8066 | 0.8382 | 0.7729 | 0.7749 |

The increase in SNR can be seen qualitatively in representative spectra taken from two voxels in a healthy subject (Figure 2). It can be seen that the spectra reconstructed using U-Net2 are less noisy than those reconstructed using MLP. Spectra reconstructed using U-Net2 are also qualitatively most similar to the spectra from the fully sampled data taken from the same region across all acceleration factors and spectra from the 4× accelerated data are less noisy than the spectra from the 6× and 7× accelerated data.

**Figure 2.**
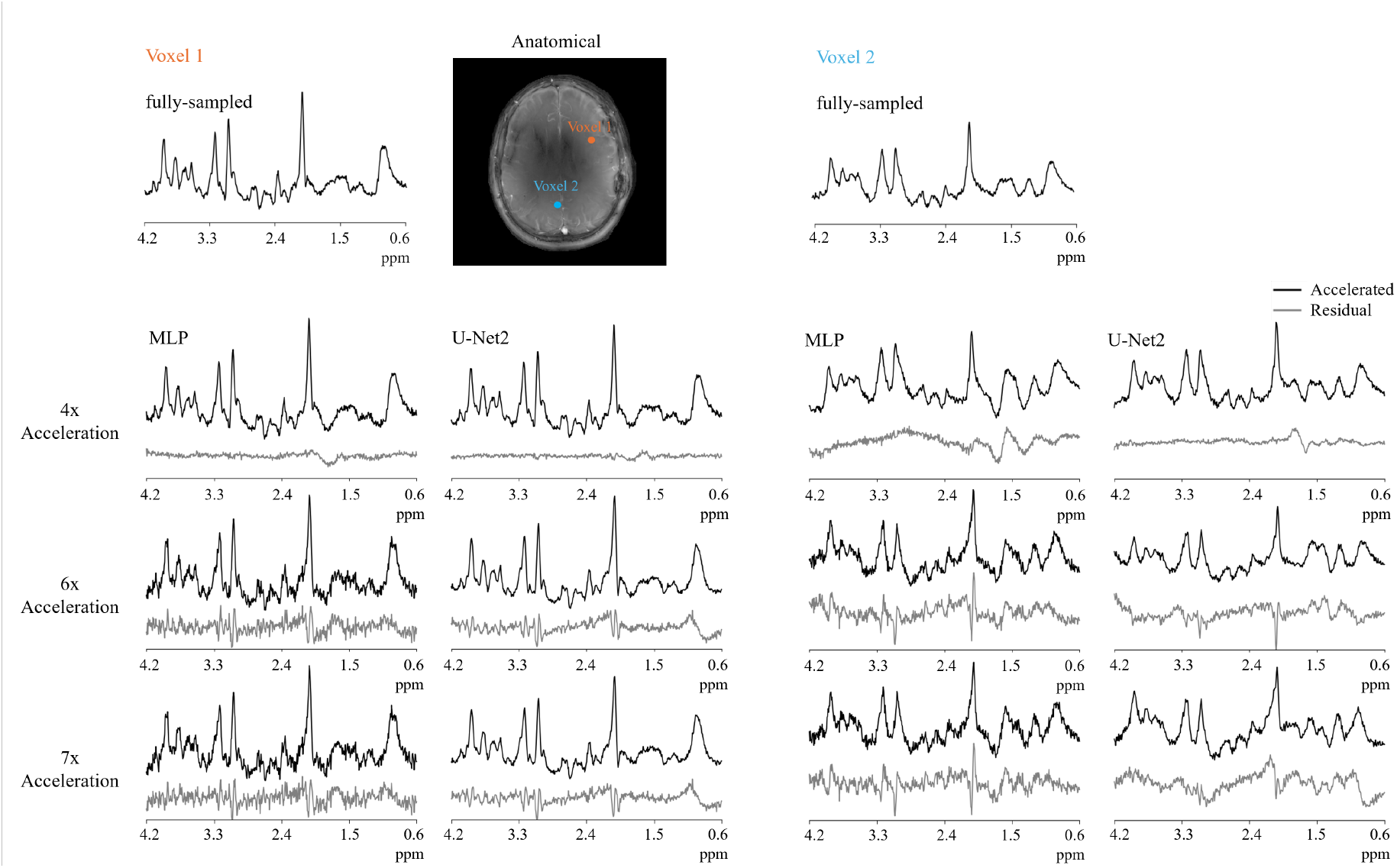
Representative accelerated and non-accelerated spectra taken from two voxels of ^1^H MRSI data acquired in one healthy subject. At all acceleration rates, the spectra reconstructed with MLP are visibly noisier than that reconstructed with U-Net2. In all voxels, the spectra taken from the fully sampled data are qualitatively similar to the accelerated spectra taken from the same region.

Figure 3 and Figure 4 present metabolite maps for Cr, NAA, Cho, Glx and mI for fully sampled, 4× accelerated, 6× accelerated and 7× accelerated data reconstructed with MLP and U-Net2 models from one healthy volunteer (Figure 3) and one brain tumor patient (tumor location: L Frontal Temporal) (Figure 4) respectively. As expected, maps obtained at lower acceleration rates (4×) exhibit greater similarity to the fully sampled reference, aligning with the SSIM index results. U-Net reconstructions consistently demonstrate smoother distributions in Cr, NAA, and Glx at low acceleration rate.

**Figure 3.**
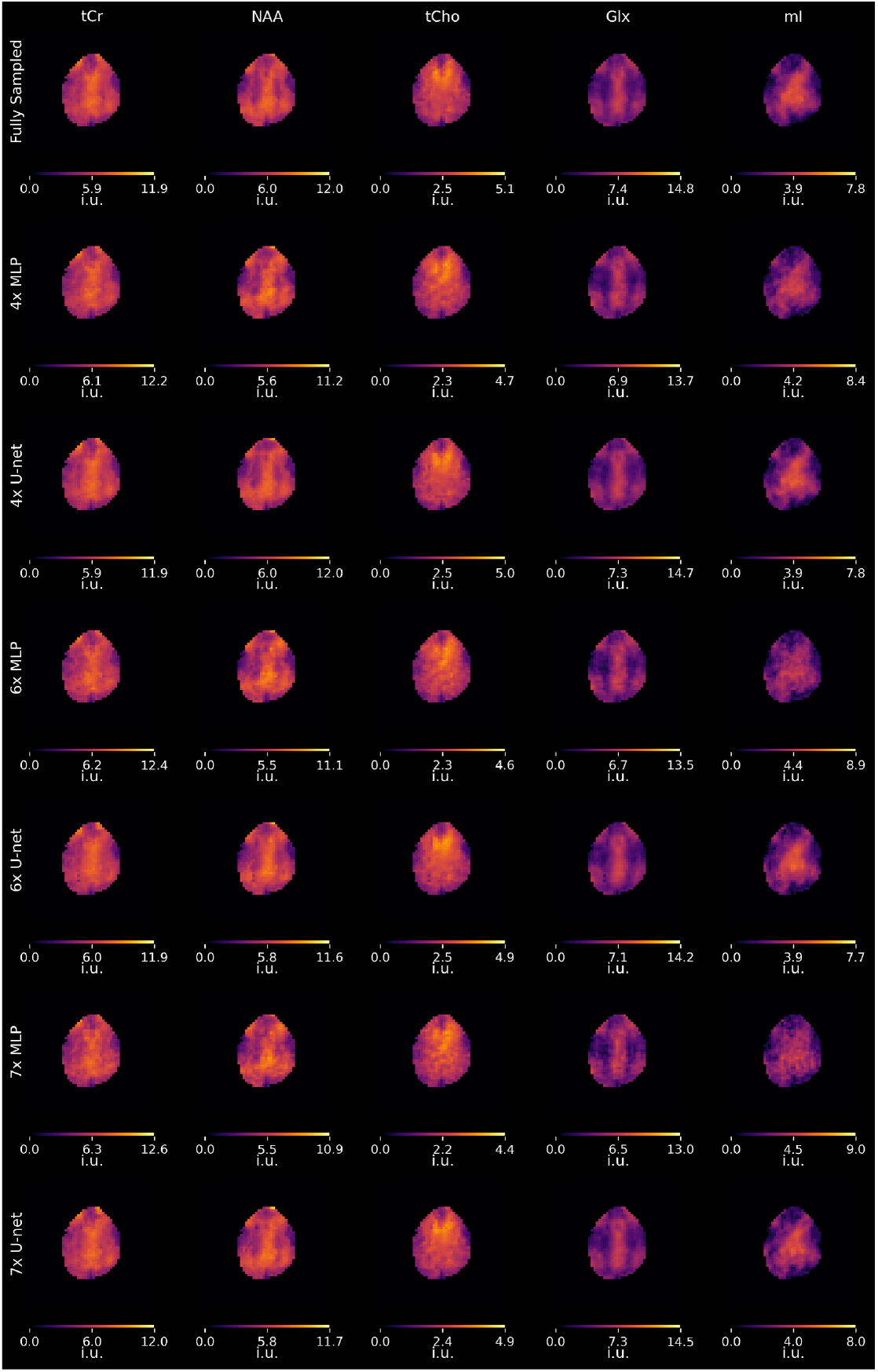
Cr, NAA, Cho, Glx and mI metabolic maps in a healthy volunteer for the fully sampled k-space data, 4× accelerated data, 6× accelerated data and 7× accelerated data reconstructed using MLP and U-Net2.

**Figure 4.**
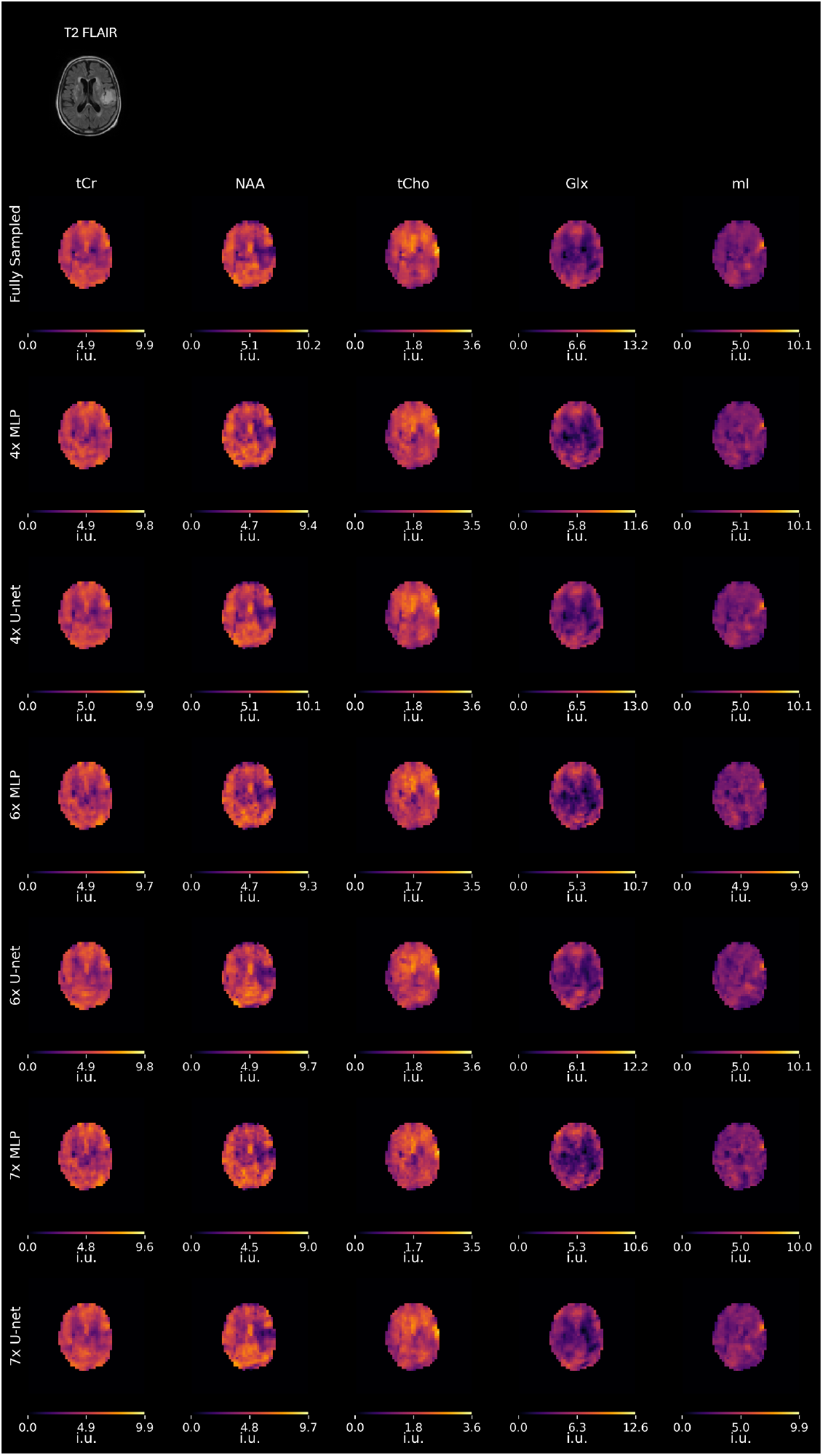
Cr, NAA, Cho, Glx and mI metabolic maps in a patient with low-grade glioma for the fully sampled k-space data, 4× accelerated data, 6× accelerated data and 7× accelerated data reconstructed using MLP and U-Net2.

## 4. Discussion

The prolonged acquisition times of ^1^H MRSI underscore the need for methods to improve scan efficiency. Optimizing acquisition speed not only reduces examination duration but also enables integration of MRSI with multimodal MRI data, amplifying the diagnostic utility of ^1^H MRSI in disease characterization. While neural-network-based GRAPPA acceleration has shown superior performance in comparison to classical parallel imaging reconstruction methods for high-SNR ^1^H MRSI at 9.4T (Nassirpour et al., 2018), semi-synthetic calibration techniques have addressed reconstruction challenges for lower-SNR ^1^H MRSI data at 7T (Chan et al., 2022). In this study, we demonstrate that applying U-Net architectures to lower-SNR ^1^H MRSI datasets further advances reconstruction quality and offers better SNR retention, offering an additional leap in performance and acceleration.

Our results demonstrate that U-Net–based models substantially enhance SNR and reduce lipid-related root mean square error (RMSE) across a range of acceleration rates. Among the tested architectures, U-Net2 achieved the greatest improvements in SNR, the most effective suppression of lipid artifacts, and the highest structural similarity. This superior performance likely arises from the model’s ability to exploit convolutional networks for robust k-space feature extraction, thereby improving the prediction of missing k-space data points. In evaluating deep learning–based reconstructions, both the acceleration factor and the network architecture are critical determinants of performance. While higher acceleration rates shorten acquisition time, they also introduce additional noise and potential information loss.

High SSIM values (≥0.91 at 4×, ≥0.78 at 6 ×, and ≥ 0.70 at 7×, averaged across the same metabolites) confirmed that the spatial distribution of metabolites was well preserved following AI reconstruction. Taken together, these findings suggest that U-Net2 achieves an optimal balance among acquisition speed, noise suppression, and structural fidelity, making it a strong candidate for clinical translation. In addition, the results highlight the importance of training strategies in supporting model generalization. Future work will focus on systematically evaluating diverse deep learning architectures within the current re-construction pipeline and optimizing computational workflows for clinical scalability.

Compared with other state-of-the-art AI-based MRSI reconstruction methods, our Cartesian U-Net approach offers distinct advantages by directly predicting missing k-space points to accelerate acquisition. The Deep-ER framework employs deep learning on non-Cartesian ECCENTRIC acquisitions, reporting substantial improvements in SNR and Cramer–Rao lower bounds(Weiser et al., 2025). In contrast, our method is designed for Cartesian sampling, which remains the dominant clinical acquisition scheme due to its straightforward calibration, compatibility with existing scanner platforms, and robustness against trajectory-related artifacts(Wright et al., 2014). Additionally, our U-Net is multimodal, leveraging T1-weighted anatomical images to guide k-space prediction, further improving reconstruction quality. Finally, building upon the conventional GRAPPA-based MultiNet framework, our method inherits reduced aliasing artifacts while achieving true acceleration via learned k-space reconstruction.

In summary, this study underscores the transformative potential of deep learning, particularly U-Net variants, in advancing MRSI reconstruction. By refining training protocols, optimizing acceleration parameters, and addressing computational bottlenecks, these methods promise to elevate diagnostic imaging quality while drastically curtailing scan times.

## Acknowledgements

This work was performed at the facilities of the Advance Imaging Research Center (AIRC) at the University of Texas Southwestern Medical Center (UTSW) and supported by funding from the Cancer Prevention and Research Institute of Texas (CPRIT) grant / RR180056.

